# Choice of medium affects PBMC quantification, cell size, and downstream respiratory analysis

**DOI:** 10.1101/2022.01.10.475633

**Authors:** Ida Bager Christensen, Lucas Ribas, Steen Larsen, Flemming Dela, Linn Gillberg

## Abstract

High-resolution respirometry (HRR) can assess PBMC bioenergetics, but no standardized medium for PBMC preparation and HRR analysis exist. Here, we study the effect of four different media (MiR05, PBS, RPMI, Plasmax) on quantification, size, and HRR analysis (Oxygraph-O2k) of intact PBMCs. Remarkably, PBMC quantification was 21% higher in MiR05 than PBS and Plasmax, and 28% higher than in RPMI, causing O_2_ flux underestimation during HRR due to inherent adjustments. Moreover, smaller cell size of PBMCs and aggregation was observed in MiR05. We suggest optimization of HRR with a standardized, plasma-like medium for future HRR analysis of intact PBMCs.

## 1. Introduction

Apart from the prominent role in energy metabolism, mitochondria are mainly known for their pronounced roles in cellular signaling, differentiation, proliferation, and apoptosis^1–3^. Mitochondrial function relies on their dynamic nature and bidirectional regulation with the cell nucleus, and these homeostatic factors are influenced by the cellular metabolic status and the surrounding microenvironment^4–6^.

Mitochondrial respiratory function is intrinsically linked to oxidative phosphorylation (OXPHOS) and ATP demands of the cell. High-resolution respirometry (HRR) provides dynamic measurements of the respiratory capacity of mitochondria by determining the oxygen consumption rate of cells and tissues^7,8^. Most HRR studies have so far focused on mitochondrial function in solid tissues such as skeletal muscle and adipose tissue. There is a great potential in HRR analysis of on circulating peripheral blood mononuclear cells (PBMCs) which meet energy metabolic requirements by OXPHOS rather than glycolysis^9^. Blood cells are exposed to a wide range of stimuli in the vasculature and can undergo drastic metabolic changes in relation to physiological and immunological homeostasis^3,10–12^. Thus, assessment of respiratory function in blood cells provide important information on immunological activity. Recently, bioenergetic alterations of PBMCs has been shown to associate with e.g., septic shock, type 2 diabetes, and major depression disorder^13–16^.

Accurate measurements of respiratory function and OXPHOS capacity of PBMCs depends on optimized, standardized preparation and execution of HRR. This secures reproducibility and enable precise data interpretation. Despite an increasing number of studies involving blood cell bioenergetics, no standards of what media to dissolve blood cells in prior to cell quantification and HRR analysis currently exist. Mitochondrial Respiration media 05 (MiR05) is specifically designed to support mitochondrial respiration for analysis of mitochondrial preparations such as isolated mitochondria, tissue, or cellular preparations^17^. RPMI 1640 medium (RPMI) is a growth medium developed for culturing lymphocytes^18^, however, its supraphysiological levels of nutrients may affect energy metabolism. Plasmax™ is a newly developed cell culture medium which mimics the physiological profile of human plasma^19,20^. Lastly, phosphate buffered saline (PBS) is widely used for biological applications. The aim of this study was to investigate the effect of the four media MiR05, PBS, RPMI, and Plasmax on cell quantification, cell size, and accurate HRR analysis of intact human PBMCs.

## 2. Materials and methods

### 2.1. Media used for dissolving of PBMCs

The following media were used in the study: Mitochondrial Respiration Media 05 (MiR05; Oroboros Instruments, Austria), Phosphate Buffered Saline (PBS; Gibco™, ThermoFisher Scientific, Waltham, MS, USA), RPMI 1640 Medium (Gibco™) and Plasmax™ (Ximbio, Cancer Research UK, London, UK). MiR05 was prepared onsite consisting of sucrose 110 mM, HEPES 20 mM, taurine 20 mM, K-lactobionate 60 mM, MgCl_2_ 3 mM, KH_2_PO_4_ 10 mM, EGTA 0.5 mM and BSA 1 g/L (pH 7.1).

### 2.2. Peripheral blood mononuclear cell isolation

Blood (16 mL) were taken from healthy, young volunteers via venous puncture in K2EDTA tubes, transferred to a 50 mL Falcon tube and diluted in PBS to a final volume of 30 mL. The diluted blood was layered on top of 15 mL Lymphoprep™ density gradient medium (StemCell Technologies Inc., Vancouver, Canada) in a 50 mL Falcon tube. The tube was centrifuged at 800 x g (acceleration 9; deceleration 2) for 15 min to separate PBMCs from granulocytes and erythrocytes. PBMCs were carefully collected with a Pasteur pipette and transferred to a separate 50 mL Falcon tube. The isolated PBMCs were diluted with PBS, thoroughly mixed, and evenly distributed to four new Falcon tubes. These tubes were centrifuged at 433 x g and the supernatant was discarded. PBMC pellets were resuspended in 1 mL of MiR05, PBS, RPMI or Plasmax, respectively.

### 2.3. Peripheral blood mononuclear cell quantification

From each of the four tubes with dissolved PBMCs, 10 μL was transferred to an Eppendorf tube containing 90 μL PBS. Next, 5 μL Solution 13 AO-DAPI (ChemoMetec, Lillerød, Denmark) was added, the sample shortly vortexed, and 10 μL of the solution loaded onto a NC-Slide A8™ (ChemoMetec). For quantification of live PBMCs, the Viability and Cell Count Assay on a

NucleoCounter^®^ NC-3000™ automated system using ChemoMetec NucleoView NC-3000 software (ChemoMetec) was used. The CASY^®^ Cell Counter (Roche Innovatis AG) was used for technical validation of total PBMC quantification (cell size > 6 μm) for three experiments.

### 2.4. Osmolality measurements and adjustment

To investigate if the high osmolality of MiR05 (315 mOsm/kg) affected the PBMCs, the osmolality of MiR05 was adjusted to be within the human plasma range (~275-295 mOsm/kg) by dilution with H_2_O. Osmolality of MiR05, the prepared MiR05 dilution, PBS, RPMI and Plasmax were measured using a K-7400S Semi-Micro Osmometer (KNAUER, Berlin, Germany).

### 2.5. Differential Interference Contrast imaging

PBMCs were isolated as described, dissolved in MiR05 at 315 mOsm/kg, MiR05 at 290 mOsm/kg, PBS, RPMI or Plasmax and left for 30 minutes for the cells to adapt to the different media. All tubes were centrifuged 3 min at 12.000 x g and the supernatant was discarded. The PBMCs were resuspended in Zamboni fixative (2%, pH 7.4) and left for one hour. Next, samples were centrifuged 30 seconds at 5000 rpm, the Zamboni fixative supernatant was discarded, and PBMCs were resuspended in PBS. The whole volume of sample was mounted to microscope slides, covered with coverslips (#1.5 thickness) and sealed with nail polish. To image the PBMCs, standard Differential Interference Contrast (DIC) imaging was performed on a confocal laser scanning microscope (Carl Zeiss, LSM 710, Oberkochen, Germany). For overview images, imaging was performed through a Plan-Apochromat 20x/0.8 objective, while higher resolution images were obtained from selected areas through an EC Plan-Neufluar 40x/1.30 Oil DIC objective. Tilling was performed with 10% overlap. Image analysis was performed using Fiji (ImageJ, v. 2.1.0/1.53 c, NIH, Bethesda, Maryland, USA). Ten cells from each tile were randomly selected and cell area was manually defined and measured. As the main selection criteria, selected cells were to be in focus and separated (not in aggregates). The area of 90 cells for each media were measured.

### 2.6. Fluorescence-activated cell sorting

For fluorescence-activated cell sorting (FACS) analysis, 100,000 PBMCs that had been dissolved and quantified in each media were stained with antibodies for CD3 (APC-H7, BD Biosciences (BD)), CD4 (Alexa Fluor^®^ 700, BD), CD8 (FITCH, BD), CD14 (PE-Cy7™, BioLegend), CD19 (APC, BD), CD56 (Super bright 600, BD), and CD45 (Pac blue, BD). 7-AAD was used as viability marker. Samples were washed in PBS, re-dissolved in the different media and run on a CytoFLEX S flow cytometer with the CytExpert 2.4 software (Beckman coulter, Indianapolis, IN, USA). The results were analyzed in FlowJo v. 10.8.0 (Becton, Dickinson and Company, Ashland, OR, USA).

### 2.7. High-resolution respirometry

Respiratory capacity of intact PBMCs was measured by high-resolution respirometry (Oxygraph-O2k, Oroboros Instruments, Innsbruck, Austria) at 37°C and 750 rpm stirring. Data was recorded by DatLab 6.1.0.7 software (Oroboros Instruments). Instrumental background calibration was routinely performed according to the manufacturer’s guidelines. All experiments were performed in 2 mL chambers containing MiR05. Oxygen calibration at air saturation was performed immediately before each experiment. Calculations of O_2_ flux per million PBMCs were based on the specific barometric pressure, oxygen concentration in the chamber and the oxygen solubility factor of MiR05 (0.92). Experiments were performed at an oxygen concentration ranging from 0 to 200 μM. Based on cell counts for PBMCs in the different media, a cell suspension volume corresponding to 1.5 million PBMCs was determined. This volume of MiR05 was removed from the chambers, where after the same volume of cell suspension was added. Then, basal O_2_ flux of the PBMCs was measured.

### 2.8. Statistical analyses

All statistical analyses were performed in RStudio (v. 1.3.1093). Shapiro-Wilk Normality Tests were performed to test for Gaussian distribution. F Tests were performed to test for equal variance of PBMCs dissolved in the different media. Statistical testing of cell quantification and basal respiration was performed as paired analyses, and statistical significance was determined using Permutation *t*-tests due to a small sample size, with variance equality as determined by preceding F tests. Statistical testing of cell sizes was performed as unpaired analyses, and statistical significance was determined using Student’s *t*-tests with variance equality as determined by preceding F tests. A significance threshold of *α* = 0.05 was used for all statistical tests.

## 3. Results

### 3.1. Type of media affects PBMC quantification

First, we investigated if there was any difference in cell count when PBMCs were dissolved in the different media (MiR05, PBS, RPMI and Plasmax) and quantified on a NucleoCounter^®^ NC-3000™. Ten experiments investigated the effect of PBMCs dissolved in MiR05, PBS or Plasmax; nine experiments furthermore included RPMI. Interestingly, eight of ten experiments showed higher cell quantification of PBMCs dissolved in MiR05 compared to PBS. Cell counts were 17–62% higher for PBMCs dissolved in MiR05 compared to PBMCs dissolved in PBS in these eight experiments (**Figure 1; Table 1**; *p* = 0.01;). Cell quantification of PBMCs in MiR05 tended to be higher than PBMCs in RPMI (**Table 1**; *p* = 0.11) or Plasmax (**Table 1**; *p* = 0.06). In contrast, cell counts of PBMC in PBS were not different from RPMI or Plasmax. Quantification of PBMCs in RPMI and Plasmax were also comparable (**Table 1**). Overall, the mean quantification ratio of PBMCs dissolved in MiR05 compared to PBMCs dissolved in PBS, RPMI or Plasmax was 21–28% higher (**Table 1**). To investigate variability of cell counts within each medium, three replicates per medium were measured for selected experiments. There were no distinct differences in variability for repeated cell count measurements for PBMCs dissolved in the different media (**Figure S1**).

**Figure 1.**
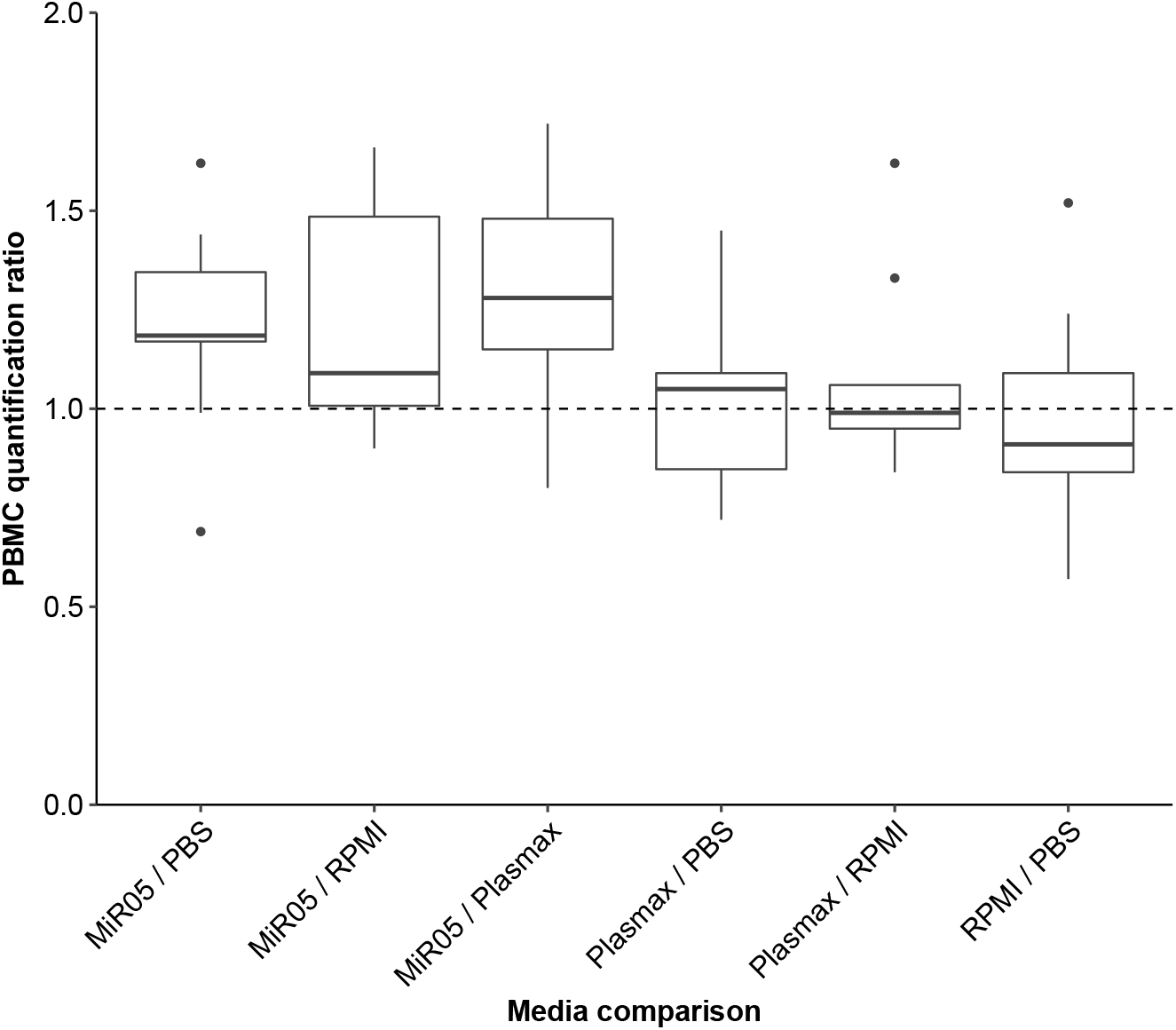
Ratios of cell counts for PBMCs dissolved in different media. Data are represented by a boxplot with whiskers indicating medians ± interquartile range (25^th^ to 75^th^ percentile). The data points represent individual experiments (Experiment 1-10). Transparency is applied to show overlapping points. The dashed horizontal line reflects identical quantification of live PBMCs dissolved in the two different media (ratio equal to 1). Ratios of PBS/RPMI, PBS/Plasmax and RPMI/Plasmax are not shown, as they are reciprocal to ratios presented.

**Table 1.** Relative quantification of PBMCs dissolved in different media. PBMCs from ten experiments were dissolved in either MiR05, PBS, RPMI or Plasmax and quantified by a Nucleo-Counter^®^ NC-3000™. Results are presented as mean quantification ratios of all ten experiments relative to a reference medium (stated above column). Statistically significant difference in live PBMC quantification for MiR05 relative to PBS, RPMI and Plasmax is indicated by * (p<0.05) and a tendency towards significance by # (p<0.10).

|  | Relative to PBS |  |  | Relative to RPMI |  |  | Relative to Plasmax |  |  |
| --- | --- | --- | --- | --- | --- | --- | --- | --- | --- |
|  | MiR05 | RPMI | Plasmax | MiR05 | PBS | Plasmax | MiR05 | PBS | RPMI |
| **Mean quantification ratio** | 1.21* | 0.98 | 1.02 | 1.28<sup>#</sup> | 1.09 | 1.07 | 1.21<sup>#</sup> | 1.02 | 0.97 |
| **Coefficient of Variation (%)** | 19.8 | 26.4 | 21.3 | 22.7 | 28.6 | 23.3 | 23.5 | 22.1 | 18.6 |

To validate the findings of varying cell counts between the media using the NucleoCounter^®^, PBMCs from three randomly selected experiments were additionally counted using a CASY^®^ Cell Counter. Like the NucleoCounter^®^, CASY^®^ revealed higher total cell counts of PBMCs dissolved in MiR05 compared to PBS, RPMI or Plasmax (**Table S1**). For one experiment, the PBMC count in MiR05 relative to PBS, RPMI and Plasmax varied distinctly between CASY^®^ and NucleoCounter^®^ measurements. However, in general the results showed comparable PBMC quantification ratios for the two methods with on average 33-67% higher cell count ratios for cells dissolved in MiR05 compared to any other media (**Table S1**). These results technically validate the NucleoCounter^®^ results and suggest that MiR05 affects PBMCs in a way which has an impact on the quantification of PBMCs using both fluorescent as well as non-fluorescent quantification techniques.

### 3.2. MiR05 affects PBMC size and aggregation propensity

Due to the dissimilarity in cell quantification observed when PBMCs were dissolved in MiR05 compared to the other media, we hypothesized that physical parameters of the PBMCs could be affected. Because osmolality can affect cell size, we measured osmolality of all media. Osmolality was high for MiR05 (315 mOsm/kg) and in the human plasma range (275–295 mOsm/kg) for PBS (296 mOsm/kg), RPMI (296 mOsm/kg) and Plasmax (287 mOsm/kg). To see if the higher osmolality of MiR05 affected the cell size, the osmolality of MiR05 was adjusted to 290 mOsm/kg by dilution with H_2_O, for further investigations of cell size. Next, Differential Interference Contrast imaging (DIC) was used to visualize PBMCs in MiR05 at 315 mOsm/kg, MiR05 at 290 mOsm/kg, PBS, RMPI and Plasmax, and the mean cell size area of PBMCs in each media was determined (**Table S2**).

Interestingly, the mean cell size areas measured in the four media were significantly different for all comparisons (all *p*-values < 0.001) except two (**Figure 2**). The mean cell size area of PBMCs dissolved in MiR05 at 315 mOsm/kg and Plasmax were similar (*p* = 0.45), and PBMCs in MiR05 at 315 mOsm/kg had tendency to be smaller than when dissolved in MiR05 at 290 mOsm/kg (*p* = 0.12). Besides variation in cell size, overview DIC images furthermore showed presence of large electrodense areas in MiR05 samples, suggesting increased aggregation propensity of PBMCs resuspended in MiR05 at both 315 and 290 mOsm/kg (**Figure S2A**). Larger aggregates were observed for PBMCs dissolved in MiR05 at 315 mOsm/kg compared to MiR05 at 290 mOsm/kg (**Figure S2A-B**). Electrodense areas were not observed in samples of PBMCs dissolved in PBS, RPMI or Plasmax (**Figure S2C-E**). No cell specific fluorescent staining was used to distinguish different cell populations of the isolated PBMCs, why it was not possible to determine the PBMC cell type in the aggregates. The distribution of various cell areas can be explained by the cell population consisting of both T-lymphocytes, B-lymphocytes, NK cells and monocytes.

**Figure 2.**
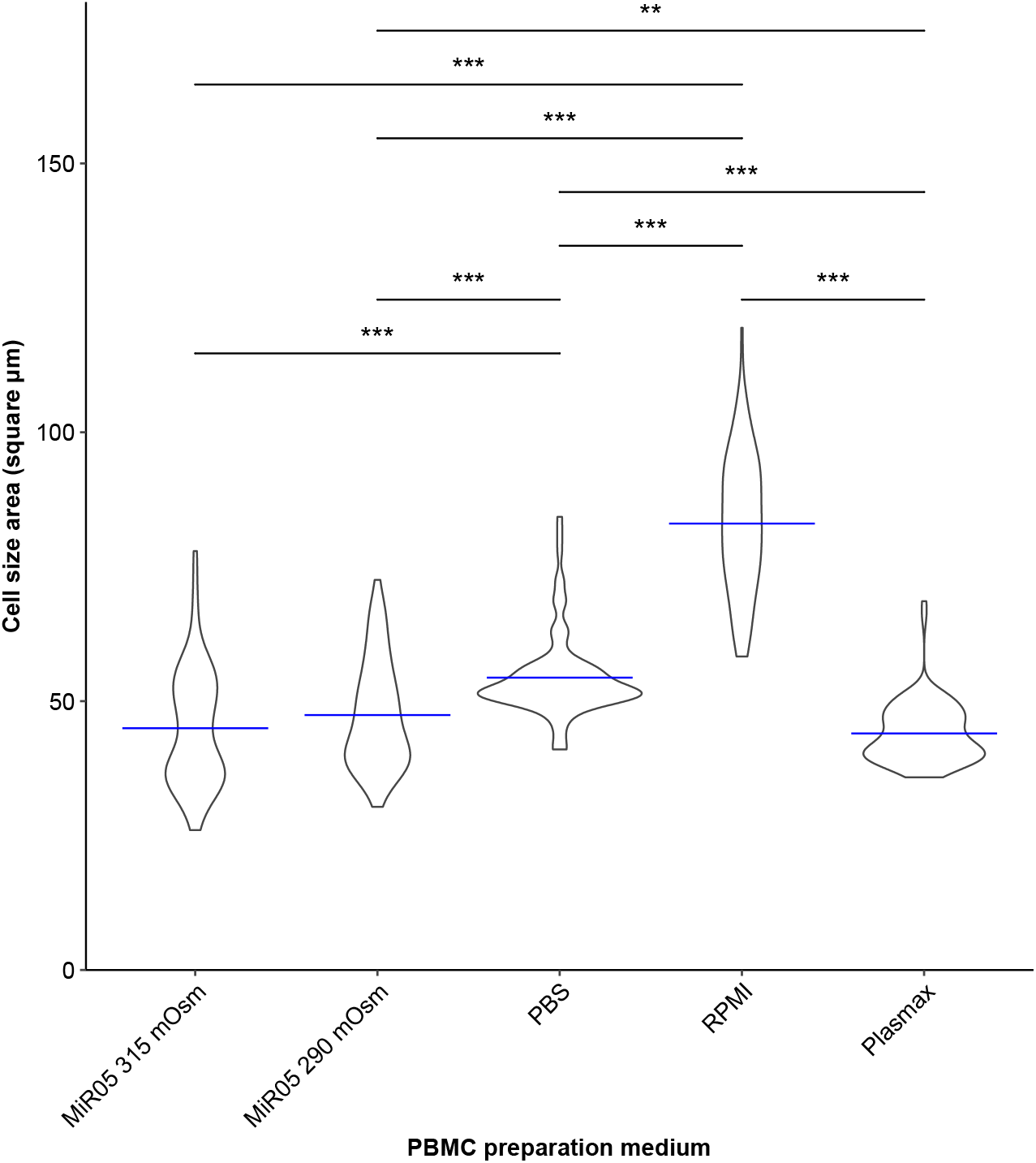
Cell size area of PBMCs dissolved in the different media. Violin plot showing the distribution of the quantitative measurements of PBMC size area after being dissolved in MiR05 at 315 mOsm/kg, MiR05 at 290 mOsm/kg, PBS, RPMI and Plasmax. Cell size area of 90 PBMCs dissolved in each medium were measured. The cell size areas were measured in DIC images (LSM 710) using Fiji, ImageJ. Blue lines mark mean cell size area. All comparisons are significantly different except MiR05 315 vs. 290 mOsm/kg and MiR05 315 mOsm/kg vs. Plasmax. ***: p-value < 0.001, **: p-value < 0.005.

### 3.3. FACS analysis supports altered cell size and aggregation propensity in MiR05

To further evaluate how MiR05 (315 and 290 mOsm/kg), PBS, RPMI and Plasmax affected the physical parameters of PBMCs, the PBMCs were stained with antibodies for all leukocytes (CD45+) and for the different types of PBMCs (total CD3+ T-cells, CD4+ helper T-cells, CD8+ cytotoxic T-cells, CD14+ monocytes, CD19+ B-cells and CD56+ NK-cells), run on a CytoFLEX S Flow Cytometer and analyzed in FlowJo. First, we gated the presumed PBMCs on a forward scatter and side scatter (FSC/SSC) histogram and found that PBMCs dissolved in MiR05 (315 and 290 mOsm/kg) had a drop in forward scatter for the main proportion of cells, which indicate a drop in cell size, compared to PBMCs dissolved in the other media (**Figure 3**). When gates for the different cell types were set up, it was obvious that cell counts in gates for cytotoxic T-cells (CD8+) and monocytes (CD14+) were drastically lower when dissolved in MiR05 compared to PBS, RPMI or Plasmax (**Table S3**; **Figure S3**). Instead, we observed a population of cells with lower forward scatter, indicating smaller cell size, that were not within any of the gates for the different cell types, in the MiR05 dissolved samples. The total PBMC count was 33% higher and the alive, non-aggregated (singlet) PBMC count 21% higher for PBMCs dissolved in MiR05 at 315 mOsm/kg compared to PBMCs dissolved in RPMI. The highest PBMC count was however observed for PBMCs dissolved in PBS whereas the most aggregation (% non-singlets) were observed for PBMCs dissolved in MiR05 (**Table S4**). Notable, the total number of counts/events were several folds lower for MiR05 dissolved cells, making the frequency of parent (PBMC counts per total counts) three times higher for MiR05-dissolved compared to RPMI-dissolved PBMCs and five times higher than PBS-dissolved PBMCs (**Table S4**). To figure out if the frequency of parent in MiR05-dissolved samples were caused by the foamy character of MiR05, the media were run on the flow cytometer without PBMCs or antibody staining. Here, small aggregates (most likely BSA aggregates) were obvious in MiR05, but the counts disappeared in the gates for PBMCs. To sum up, FACS analysis confirmed the smaller cell size and aggregation propensity of PBMCs dissolved in MiR05 which was observed with DIC-imaging. Especially, MiR05 seem to have a drastically effect on the size of cytotoxic T-cells and the internal complexity of monocytes. Together, the multiple influences that MiR05 have on PBMCs most likely interfere with reliable quantification.

**Figure 3.**
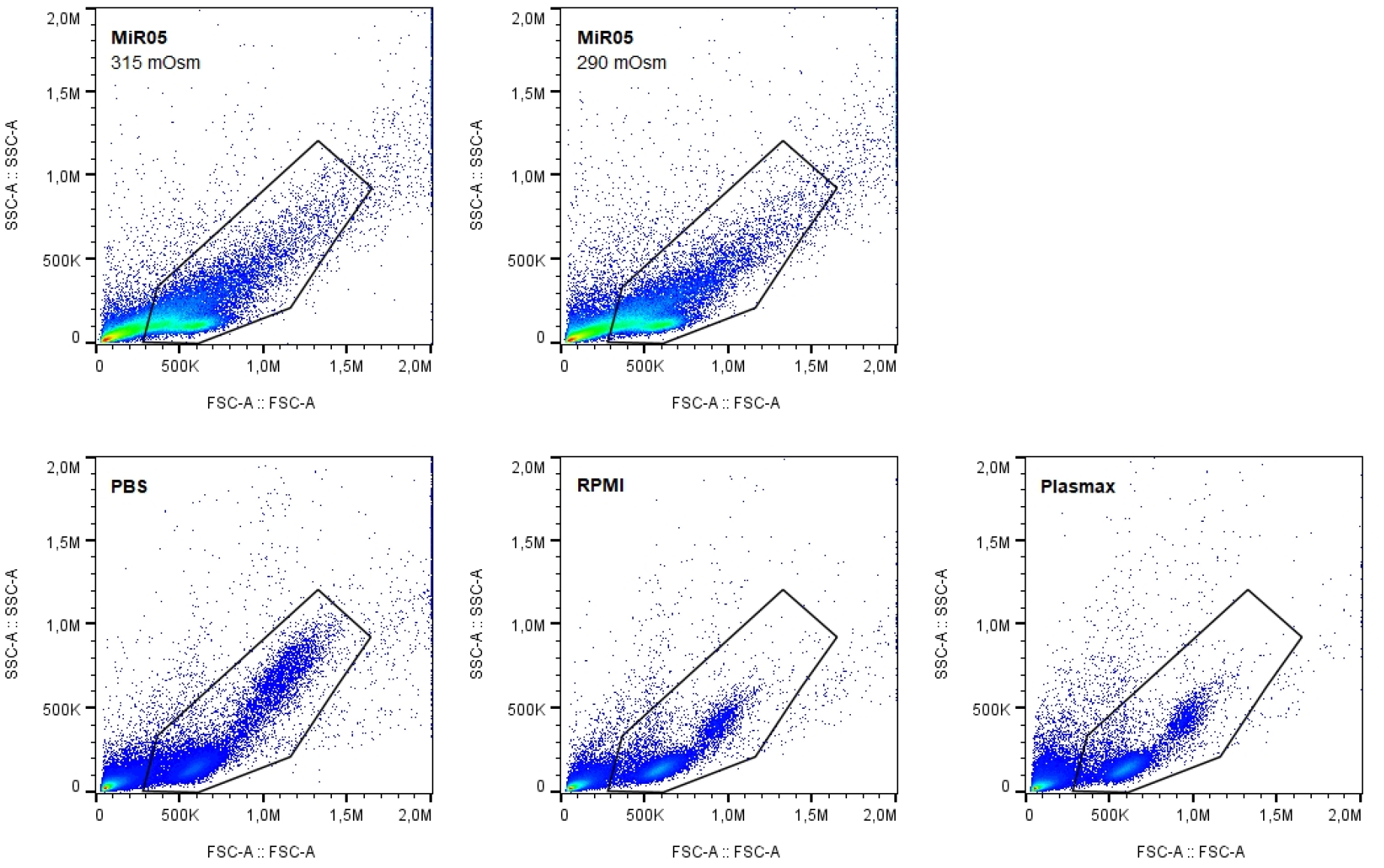
Fluorescence-activated cell sorting of PBMCs dissolved in the different media. FACS analysis performed using CytoFLEX S. The gates mark presumed PBMCs. PBMCs were dissolved in either MiR05 at 315 mOsm/kg, MiR05 at 290 mOsm/kg, PBS, RPMI or Plasmax.

### 3.4. Respiratory changes of PBMCs prepared differently is explained by cell count discrepancy

Continuous oxygen consumption measurements and the derivate O_2_ flux is adjusted for the amount of cells in the O2k chamber, which depend on accurate cell quantification. Motivated by the discrepancy of cell count, cell size, and aggregation of PBMCs observed in MiR05 samples, we investigated how basal respiration of PBMCs measured by HRR was affected by technical adjustment for 1) 1.5 million PBMCs based on cell counts performed in each medium, and 2) varying amounts of PBMCs based on the cell count of one designated medium. Because equivalent amounts of PBMCs were isolated and dissolved in different media, we hypothesized that 1) basal O_2_ flux is dependent on accurate cell count; and 2) that the media used for dissolving PBMCs (and diluting MiR05 in the O2k chamber) might affect respiration differently. Eight experiments tested this, five included RPMI. Initially, basal respiration of 1.5 million PBMCs based on cell counts performed in each of the four media were compared. Here, PBMCs dissolved in MiR05 showed a tendency towards lower basal O_2_ flux compared to PBMCs dissolved in PBS (14.7%; **Figure 4**; *p* = 0.07). Basal O_2_ flux of PBMCs dissolved in RPMI and Plasmax was not significantly different from PBMCs dissolved in MiR05 (**Figure 4**).

**Figure 4.**
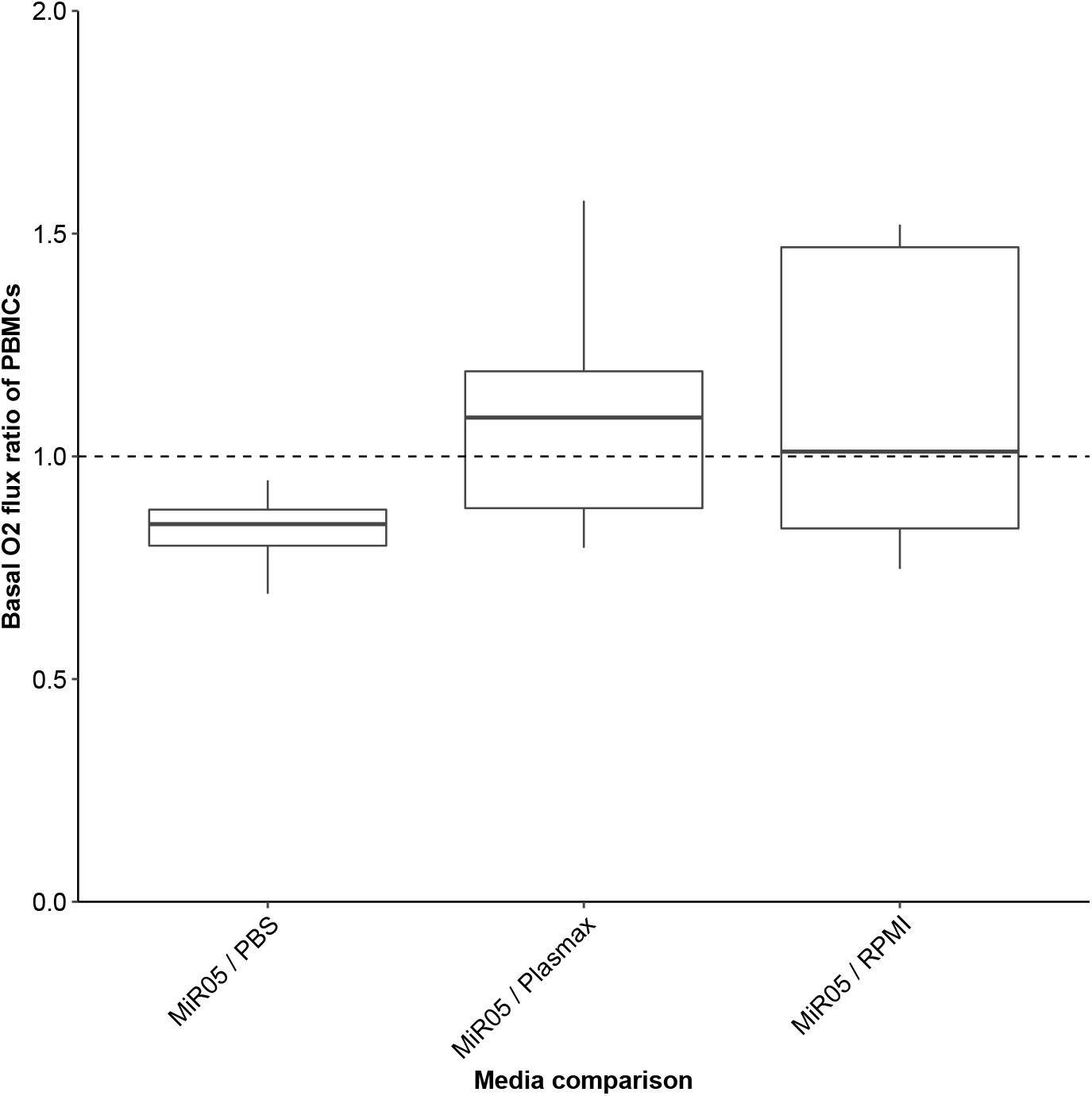
The influence of cell quantification on basal O_2_ flux of intact PBMCs measured by HRR. Basal O_2_ flux of 1.5 million intact PBMCs dissolved in MiR05, PBS, Plasmax or RPMI was measured, and O2k oxygraphs were adjusted for 1.5 million PBMCs based on cell quantification performed in each specific media. Data are represented by a boxplot with whiskers indicating medians ± interquartile range (25^th^ to 75^th^ percentile). The data points represent individual experiments (eight experiments, five for RPMI). Transparency is applied to show overlapping points.

Due to the higher cell count of PBMCs dissolved in MiR05, the volume of cell suspension added to the O2k chambers was reduced compared to PBMCs dissolved in the other media when analysing respiration of 1.5 million PBMCs. Based on this observation, we adjusted the HRR analyses in DatLab so that the amount of cells added to the chamber was based on a designated cell quantification performed in either MiR05, PBS, RPMI or Plasmax. When doing so, the amount of PBMCs changed markedly, ranging from 0.9 to 2.6 million. PBMCs dissolved in MiR05 now had similar or slightly higher basal O_2_ flux compared to PBMCs dissolved in any of the other media (**Figure 5**). This was in clear contrast to the results where O2k oxygraphs had been adjusted for cell count from each of the media, where there was a tendency towards significantly lower basal O_2_ flux in MiR05 dissolved samples (**Figure 5**; Individual cell counts; *p* = 0.07). To sum up, it is important to dissolve and quantify cells in the same medium for all samples to accurately quantify mitochondrial respiration of PBMCs using HRR.

**Figure 5.**
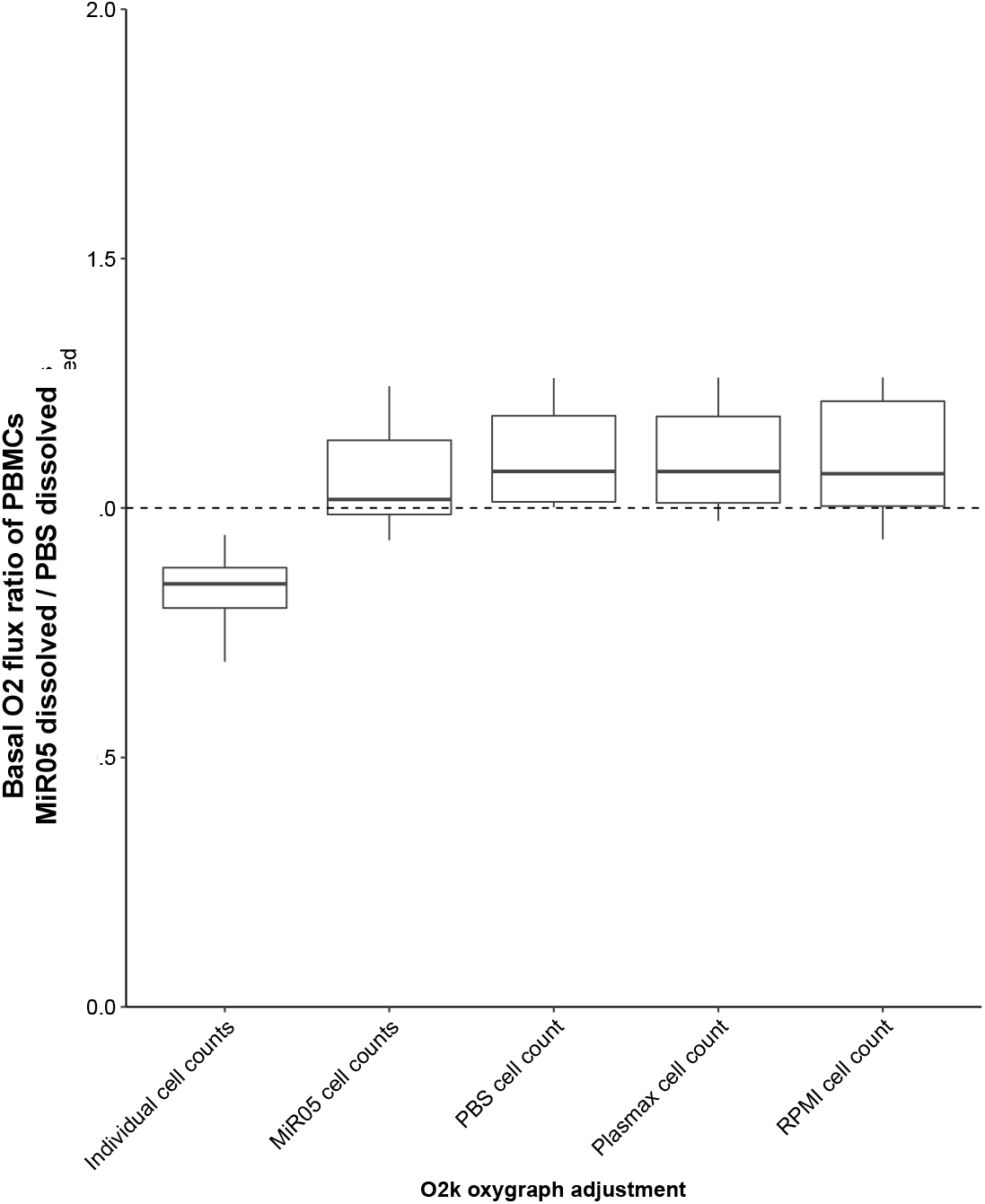
The effect of media on HRR analysis diminishes if adjusted for cell quantification performed in one media. Basal O_2_ flux ratio of intact PBMCs dissolved in MiR05 relative to PBMCs dissolved in PBS. Data are represented by a boxplot with whiskers indicating medians ± interquartile range (25^th^ to 75^th^ percentile). The data points represent individual experiments (eight experiments, five for RPMI). Transparency is applied to show overlapping points. First boxplot indicates basal O_2_ flux adjusted for PBMC quantification in MiR05 and PBS (similar to Figure 4). The following boxplots indicate when O2k-oxygraphs were adjusted for one designated cell count (i.e., independent of what medium the PBMC was dissolved in during preparation), resulting in varying number of PBMCs examined.

## 4. Discussion

In this study, we have systematically investigated how PBMCs dissolved in the media MiR05, PBS, RPMI and Plasmax affects cell quantification, aggregation propensity and respiratory flux of intact PBMCs. Using both fluorescent and non-fluorescent cell quantification, we observed a significantly higher cell count of PBMCs dissolved in the standard respiration medium MiR05 compared to PBMCs dissolved in PBS, RPMI or Plasmax. Since O2k oxygraphs are technically adjusted for the amount of cells in each chamber, the increased cell count in MiR05 caused an underestimation of O_2_ flux during subsequent HRR analysis. The overestimation of cell amount in MiR05 might be related to the physio-chemical properties of the medium like the high osmolality, which can lead to a decreased size and pronounced aggregation when PBMCs are dissolved in MiR05 as seen in this study.

The NucleoCounter^®^ NC-3000™ system is based on fluorescent cytometry where viable and non-viable cells are stained with membrane-permeable acridine orange (AO) and membrane-impermeable DAPI dyes, respectively. If PBMCs exist in large aggregates, DAPI is potentially hindered to reach cells centered in the aggregate, while AO would be unhindered due to its membrane-permeable properties. This could underestimate the number of dead cells without affecting the total cell count, resulting in an overall overestimation of viable cells in a sample. Quantification by CASY^®^ also displayed higher cell counts in MiR05 samples. Thus, two different cell quantification methods based on different techniques support the observed overestimation of PBMCs in MiR05. Interestingly, aggregation-prone cells, e.g., breast cancer cell lines, are known to be challenging for cell counting, as superimposed cells are hard to distinguish and count accurately for most automated cell counters^21^. Thus, if PBMCs are superimposed in an aggregate due to the altered physico-chemical properties of MiR05, this could explain the higher and potentially inaccurate PBMC counts. One of these properties might be the high osmolality of MiR05. Even though DIC imaging showed signs of PBMCs being slightly (but non-significantly) larger and with smaller aggregates compared to when the osmolality of MiR05 was within the plasma range (290 mOsm/kg vs 315 mOsm/kg), the PBMCs in MiR05 at both osmolalities showed similar physico-chemical properties on FACS analysis (such as distinct changes in appearance of monocytes and CD8+ T-cells), as well as aggregation propensity and decreased cell area compared to the other media as seen by DIC imaging. This indicates that other properties influenced the quantitative deviations in cell counts observed for MiR05.

Despite MiR05s effect on PBMC counts, we did not observe an increased coefficient of variation of MiR05 dissolved cells compared to other media between the ten different experiments, and we moreover did not observe increased standard deviation for repeated cell count measurements in MiR05 of a single experiment compared to the other media. Thus, our data indicate that MiR05 dissolved PBMCs has an impact on accuracy but not precision of cell count measurements.

When our HRR analyses were adjusted for the cell count performed in only one of the designated media, the basal O_2_ flux ratio of MiR05 dissolved PBMCs relative to PBS, RPMI, or Plasmax dissolved PBMCs was not equal to one. We hypothesize that the slight increase in basal O_2_ flux as observed for MiR05 dissolved PBMCs might be due to the O2k chamber being diluted with another media than MiR05 when the PBMCs were added to the chamber. The composition of the media in the chamber might affect both the cellular energy metabolism and the technical sensitivity such as the oxygen solubility of the media. This is in line with our observation that the largest difference in basal respiration of MiR05 relative to PBS dissolved PBMCs was seen in experiments where the cell suspension had a low concentration of cells, and consequently a large volume of PBS cell suspension was added to the chambers to be able to achieve 1.5 million cells. Thus, dilution of MiR05 with other media in the O2k chambers may affect respiration measurements and should be avoided. Sumbalova et al. state that choice of medium and its composition is an important parameter when evaluating respiration of living cells ^22^.

Based on the results from this study, we conclude that MiR05 cause decreased size and increased aggregation propensity of PBMCs, and overestimation of cell count (but not variability) based on both fluorescent and non-fluorescent quantification. First, we recommend to consistently use one medium for dissolving PBMCs prior to cell quantification and HRR analysis, to be able to compare inter-study respirometry data. Optimally, this media should also be used in O2k chambers during HRR analysis. Currently, MiR05 is the preferred medium for HRR analyses as it optimally supports respiration of isolated mitochondrial, tissues and permeabilized cells^17^. However, this media does not prove to be optimal for intact PBMCs based on the significant effects MiR05 has on quantification of the cells shown in this study. Moreover, we cannot rule out that the effect MiR05 has on cell size (partly due to high osmolality) and aggregation also affects oxidative energy metabolism and other cellular functions. In line with this, another media would be preferable in the future. Due to supraphysiological nutrient levels, RPMI would not be optimal for HRR analyses since this method is also used for permeabilized cells and in such respect, nutrients will provide external support for respiration and thereby interfere with the results^22^. Plasmax, which resembles human plasma, showed suitable for cell quantification, and showed no aggregation why we argue that Plasmax could be an optimal medium for HRR analysis of PBMCs. For this medium to be applied in HRR, an oxygen solubility factor, which is used for adjustment of correct O_2_ flux measurements using the O2k oxygraphs, must be determined.

## 5. Conclusions

Here, we investigated how different media affected intact human PBMCs for downstream HRR analysis to secure experimental reproducibility and enable accurate data interpretation. The standardized respiration media MiR05 caused decreased size of PBMCs, overestimated cell quantification based on both fluorescent and non-fluorescent methods, and showed increased aggregation propensity compared to PBS, RPMI and Plasmax. The overestimated cell quantification caused underestimation of basal O_2_ flux of intact PBMCs dissolved in MiR05, due to inherent adjustment for cell quantity. We conclude that methods for PBMC preparation should always be thoroughly documented to enable inter-study comparisons. Furthermore, we recommend standardizing the usage of Plasmax which better resembles human plasma, for dissolving PBMCs prior to cell quantification and for usage in O2k chambers during HRR analysis.

## 6. Acknowledgements

The authors gratefully acknowledge the Core Facility for Integrated Microscopy at University of Copenhagen, PhD Christophe Côme from Biotech Research & Innovation Centre and Epigenomlab, University of Copenhagen as well as Laboratory technician Kristoffer Racz, Department of Biomedical Sciences, University of Copenhagen for their support and assistance in this work. This work was supported by Danish Diabetes Academy (grant number NNF17SA0031406), Fru Astrid Thaysens Legat for Lægevidenskabelig Grundforskning and Dagmar Marshalls Fond.

