## Supplemental data for "Choice of medium affects PBMC quantification, cell size, and downstream respiratory analysis"

**Table S1. PBMC quantification measured by NucleoCounter® NC-3000™ and CASY® Cell Counter**
Cell quantification of total (live and dead) PBMCs dissolved in either PBS, MiR05, RPMI or Plasmax, presented as ratios relative to a reference cell count indicated at the top of the columns. The cell counts were performed by NucleoCounter® NC-3000™ and CASY® Cell Counter. Noticeable variation between methods is colored in red.

| | | Relative to PBS | | | Relative to RPMI | | | Relative to Plasmax | | |
|---|---|---|---|---|---|---|---|---|---|---|
| | **Method** | **MiR05** | **RPMI** | **Plasmax** | **MiR05** | **PBS** | **Plasmax** | **MiR05** | **PBS** | **RPMI** |
| Experiment 2 | NucleoCounter® | 1.15 | 1.04 | 1.06 | 1.10 | 0.96 | 1.01 | 1.09 | 0.95 | 0.99 |
| Experiment 2 | CASY® | 1.13 | 1.07 | 0.94 | 1.06 | 0.94 | 0.88 | 1.19 | 1.06 | 1.13 |
| Experiment 5 | NucleoCounter® | 1.46 | 1.04 | 0.96 | 1.41 | 0.96 | 0.92 | 1.52 | 1.04 | 1.08 |
| Experiment 5 | CASY® | 1.54 | 1.19 | 1.04 | 1.29 | 0.84 | 0.87 | 1.48 | 0.96 | 1.15 |
| Experiment 7 | NucleoCounter® | 0.67 | 0.58 | 0.74 | 1.15 | 1.71 | 1.27 | 0.90 | 1.35 | 0.79 |
| Experiment 7 | CASY® | 1.32 | 0.50 | 0.74 | 2.67 | 2.02 | 1.50 | 1.78 | 1.34 | 0.67 |

**Table S2. Measurements of PBMC size area in Differential Interference Contrast images**
Mean size of PBMCs dissolved in either PBS, MiR05 315 mOsm/kg, MiR05 290 mOsm/kg, RPMI or Plasmax analysed in light microscopy images recorded by LSM 710, EC Plan-Neufluar 40x/1.30 Oil DIC objective. Analyses were performed in Fiji (ImageJ, v. 2.1.0/1.53 c). A total of 90 PBMCs were measured in each medium. Mean cell size area is presented in $\mu m^2 \pm$ SD.

| | Osmolality (mOsm/kg) | Mean cell area ($\mu m^2$) |
|---|---|---|
| **PBS** | 296 | $54.4 \pm 7.6$ |
| **MiR05** | 315 | $45.0 \pm 10.9$ |
| **MiR05** | 290 | $47.4 \pm 9.8$ |
| **RPMI** | 296 | $83.0 \pm 12.7$ |
| **Plasmax** | 287 | $44.0 \pm 5.6$ |

**Table S3. FACS results of PBMCs dissolved in MiR05 315 mOsm/kg, MiR05 290 mOsm/kg, PBS, RPMI or Plasmax**

Measurements from FACS experiment analysing PBMCs dissolved in MiR05 315 mOsm/kg, MiR05 290 mOsm/kg, PBS, RPMI or Plasmax. CD45+ is a lymphocyte marker, CD3+ a T-lymphocyte marker, CD8+ a cytotoxic T-cell marker, CD4+ a helper T-cell marker, and CD19+ a B-lymphocyte marker. Results are presented as counts (upper panel) and frequent of Parent (lower panel).

| **Medium** | **CD45+ medium** Viable singlets | **CD45+ high** Viable singlets | **CD3+** CD45+ high viable singlets | **CD8+ CD3+** CD45+ high viable singlets | **CD4+ CD3+** CD45+ viable singlets | **CD19+** CD45+ high viable singlets | **Monocytes** CD45+ high viable singlets |
|---|---|---|---|---|---|---|---|
| **Count** | | | | | | | |
| MiR05 (315 mOsm/kg) | 8928 | 12832 | 7968 | 3 | 5331 | 1087 | 63 |
| MiR05 (290 mOsm/kg) | 9654 | 12122 | 7744 | 0 | 5203 | 1025 | 72 |
| PBS | 868 | 25833 | 15509 | 3963 | 10625 | 2306 | 2908 |
| RPMI | 381 | 17035 | 10781 | 3006 | 7179 | 1664 | 1252 |
| Plasmax | 299 | 17730 | 11357 | 3157 | 7593 | 1678 | 1112 |
| **Freq, of Parent** | | | | | | | |
| MiR05 (315 mOsm/kg) | 39.1 | 56.1 | 62.1 | 0.0 | 66.9 | 8.5 | 0.5 |
| MiR05 (290 mOsm/kg) | 42.6 | 53.5 | 63.9 | 0.0 | 67.2 | 8.5 | 0.6 |
| PBS | 3.0 | 89.3 | 60.0 | 25.6 | 68.5 | 8.9 | 11.3 |
| RPMI | 2.0 | 90.0 | 63.3 | 27.9 | 66.6 | 9.8 | 7.4 |
| Plasmax | 1.5 | 90.70 | 64.10 | 27.8 | 66.9 | 9.5 | 6.3 |

**Table S4. Counts based on FACS analysis of PBMCs dissolved in different media**

FACS analysis performed using CytoFLEX S to purify PBMC populations detected by flow cytometry. PBMCs were dissolved in either MiR05 at 315 mOsm/kg, MiR05 at 290 mOsm/kg, PBS, RPMI or Plasmax. Singlets indicate PBMCs that are not aggregated. Frequency of parent indicates percentage of PBMC counts/total counts.

| **Medium** | **Count** | **PBMCs** Count | **PBMC singlets** Count | **Alive PBMC singlets** Count | **Freq of Parent** (PBMCs/Count) | **PBMC singlets** (% of all) | **Alive singlet PBMCs** (% of all) |
|---|---|---|---|---|---|---|---|
| MiR05 (315 mOsm/kg) | 75789 | 25617 | 23530 | 22861 | 33.8% | 91.9% | 97.2% |
| MiR05 (290 mOsm/kg) | 76004 | 24565 | 22966 | 22657 | 32.3% | 93.5% | 98.7% |
| PBS | 455000 | 29552 | 29130 | 28931 | 6.5% | 98.6% | 99.3% |
| RPMI | 162660 | 19222 | 18993 | 18921 | 11.8% | 98.8% | 99.6% |
| Plasmax | 191635 | 19762 | 19623 | 19545 | 10.3% | 99.3% | 99.6% |

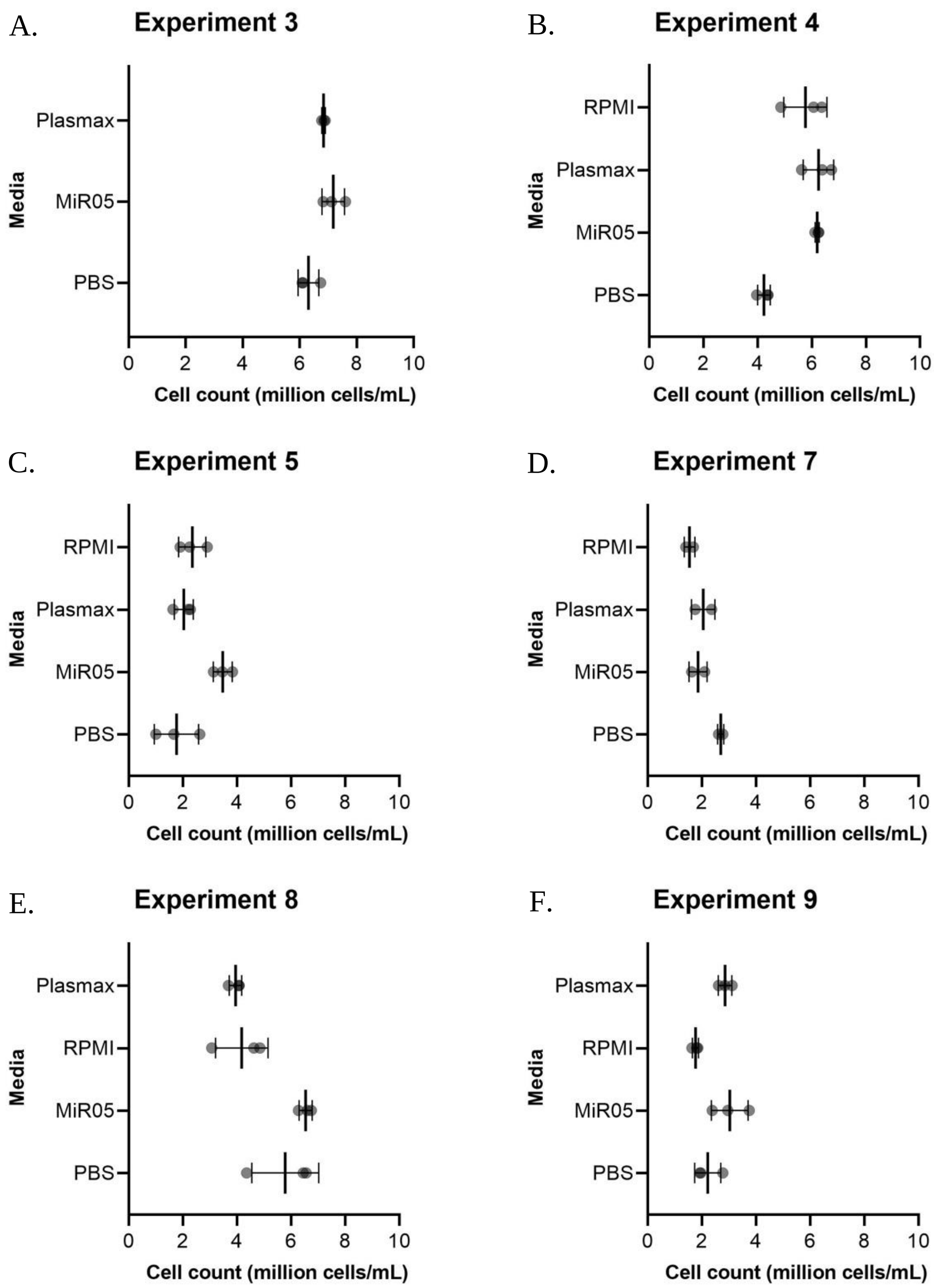

**Figure S1. NucleoCounter® NC-3000™ PBMC count replicates within individual selected experiments.** Aligned dot plot showing NucleoCounter® NC-3000™ cell count triplicates within individual experiments. A-F represents individual experiments. The data points represent cell counts triplicates performed in all four media; MiR05, PBS, RPMI and Plasmax (duplicates performed in D, Experiment 7). Mean ± SD are presented in each experiment.

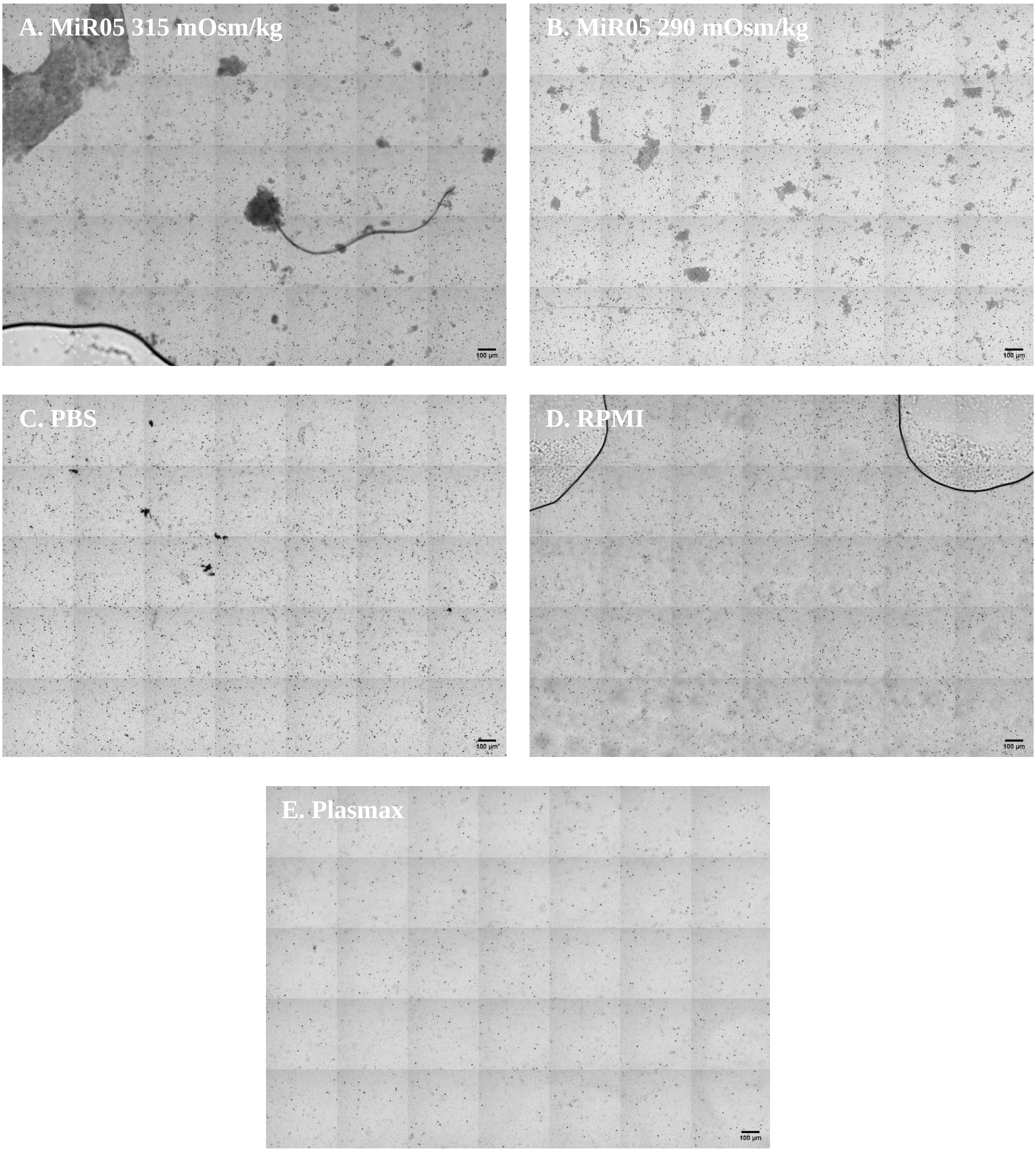

**Figure S2. Overview Differential Interference Contrast (DIC) images.**
Overview DIC images of PBMCs dissolved in different media. Microscopy was performed using Zeiss LSM 710, Plan-Apochromat 20x/0.8 objective. Scale bars show 100 µm. **A.** PBMCs dissolved in MiR05 315 mOsm/kg. Large electrodense areas are present**. B.** PBMCs dissolved in MiR05 290 mOsm/kg. Electrodense areas are present, however smaller than observed in MiR05 315 mOsm/kg. **C.** PBMCs dissolved in PBS. Electrodense areas are not present. **D.** PBMCs dissolved in RPMI. Electrodense areas are not present**. E.** Overview image of PBMCs dissolved in Plasmax. Electrodense areas are not present.

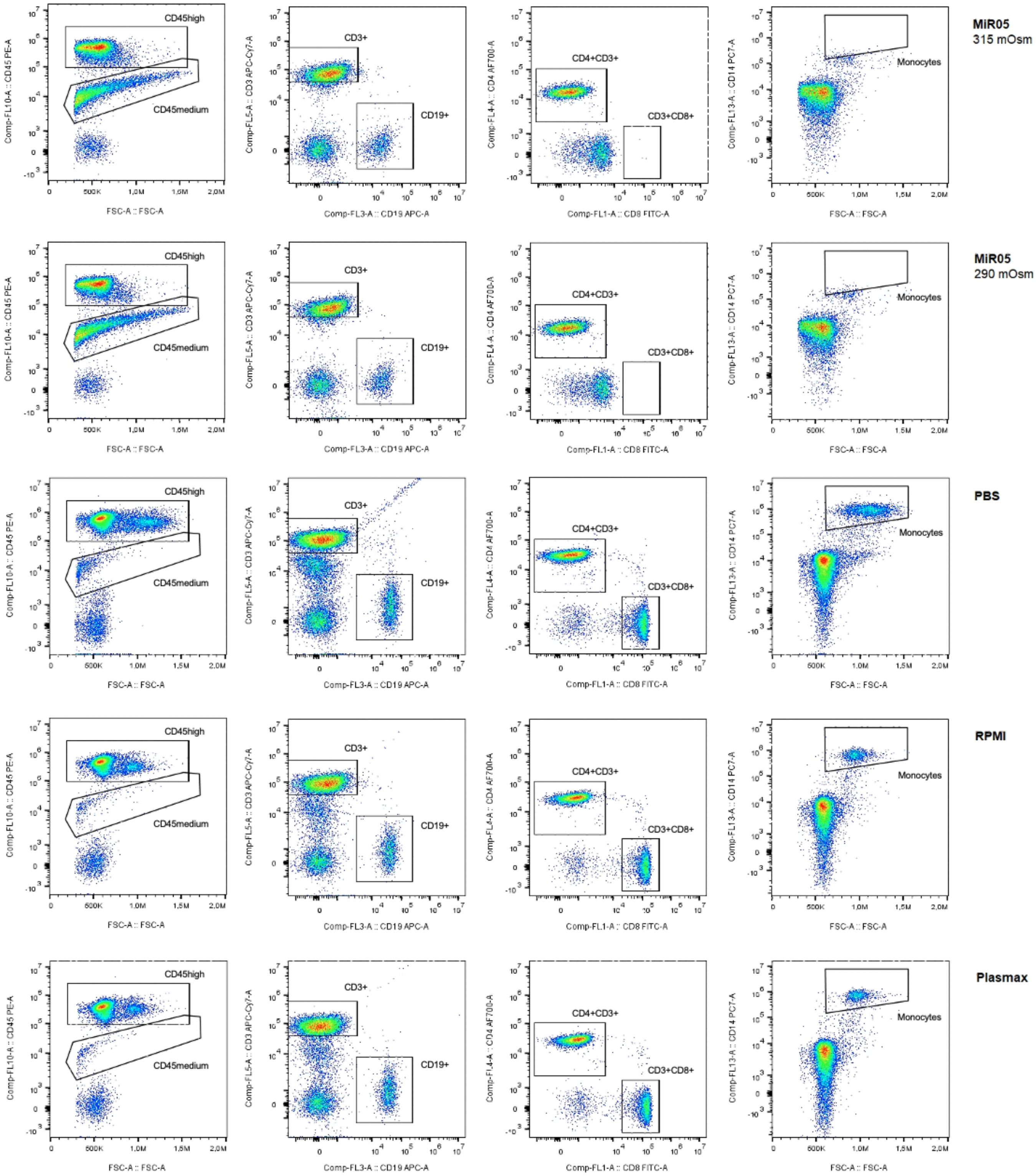

**Figure S3. Fluorescence-activated cell sorting of PBMCs dissolved in different media.**
FACS analysis performed using CytoFLEX S to purify PBMC populations detected by flow cytometry. PBMCs were dissolved in either MiR05 at 315 mOsm/kg, MiR05 at 290 mOsm/kg, PBS, RPMI or Plasmax. Gates for different cell types are marked on all graphs and cell type indicated next to the gate.
